# Circulating microRNAs as biomarkers of chronic kidney disease and its association with renal and cardiovascular outcomes in non-dialysis patients

**DOI:** 10.1101/2024.10.10.617547

**Authors:** Olalla Ramil-Gómez, Ana Cerqueira, Sara Reis Moura, Eduarda Carvalho, Núria Paulo, Isabel Brandão, Janete Santos, Patricia Quaranta, Maria João Sousa, Ana Pinho, Hugo Diniz, Maria Inês Almeida, Manuel Pestana, Inês Soares Alencastre

**Affiliations:** i3S-Instituto de Investigação e Inovação em Saúde, Porto, Portugal; Centro Hospitalar Universitário de São João, Porto, Portugal; Faculdade de Medicina da Universidade do Porto, Porto, Portugal; Endocrine, Nutritional and Metabolic Diseases Research Group, CICA-Interdisciplinary Center for Chemistry and Biology, University of A Coruña, 15071, A Coruña, Spain; Biomedical Research Institute of A Coruña (INIBIC), 15006, A Coruña, Spain; Rheumatology Research Group (GIR)-Proteomics Unit, INIBIC, 15006, A Coruña, Spain; Institute of Biomedical Sciences Abel Salazar (ICBAS), University of Porto, 4200-135, Porto, Portugal

**Author notes:** ***Corresponding author:*** Name: Inês Soares Alencastre.

**Keywords:** Nephrology, Cardiovascular disease, Non-coding RNAs, Biomarkers, Diagnosis, Prognosis

## Abstract

Chronic kidney disease (CKD) affects over 10% of the population worldwide and entails a significant risk for cardiovascular disease (CVD), leading to a 500-fold increase in cardiovascular mortality in advanced stages. Still, the pathophysiological mechanisms underlying kidney-heart intercommunication remain largely unknown. In recent years, microRNAs (miRNAs) emerged as important key regulators of gene expression that may serve as biomarkers for several diseases, however, their role in kidney-heart crosstalk in CKD remains underexplored. In this study, we evaluated the expression of a miRNA panel in plasma samples from non-dialysis CKD patients and explored their association with main comorbidities and significant outcomes in a 5-year follow-up. Results show that miR-30c-5p and miR-132-3p were downregulated in CKD patients presenting a significant power to discriminate the disease state. Importantly, only miR-30c-5p was downregulated in early glomerular filtration rate (GFR) categories, being able to discriminate these CKD patients from non-diseased individuals. Concerning cardiovascular outcomes, miR-199a-5p was found to be associated with an increased frequency of CVD. When analyzing the major disease outcomes in a 5-year time, miR-199a-5p upregulation at baseline was associated with increased mortality, while miR-324-3p was downregulated in patients who progressed to more advanced stages of the disease. These findings highlight the involvement of novel circulating miRNAs in CKD onset and progression, and identify, for the first time, the enrollment of miR-199a-5p in kidney-heart pathophysiological crosstalk, paving the way for the establishment of new biomarkers and therapeutic targets for CKD and its outcomes.

## Introduction

Chronic kidney disease (CKD) represents a growing global public health challenge and is expected to become the fifth cause of mortality by 2040 ^1^. However, the available treatments rely mainly on the implementation of healthier lifestyle habits, management of the associated comorbidities (such as hypertension or diabetes) and renal replacement therapies for advanced CKD ^2,3^. Although effective treatments to halt the progression of the disease have recently become available, namely renin-angiotensin-aldosterone system (RAAS) inhibitors and sodium-glucose cotransporter-2 (SGLT2) inhibitors, a residual risk for progression persists and, in advanced stages, CKD patients experience a reduction in life expectancy of 25 year compared to individuals with preserved renal function ^4^.

Moreover, the current markers for CKD diagnosis – GFR and albuminuria - display poor application and sensitivity for early diagnosis ^5^. In fact, although albuminuria has been associated with several CKD manifestations and CVD mortality ^6^, current worldwide rates of albuminuria screening, even in high-risk patients, are low, mainly due to albuminuria high short-term, within-person variability and the need for 24-hour measurement of albumin excretion rate (AER) for accurate assessment, that are not convenient for routine clinical practice ^6^. Moreover, although albuminuria may precede a decrease in GFR, it can be absent in tubulointerstitial or hypertensive kidney diseases and ∼30% of diabetic kidney disease patients display normal urine albumin levels ^5^. Therefore, novel diagnostic and monitoring tools are critically needed.

One of the main consequences of CKD is that it represents a significant and independent risk factor for cardiovascular disease (CVD), being the leading cause of death among CKD patients in advanced stages ^4^. Although it is known that renal and cardiac injury share common pathophysiological processes, including inflammation and fibrosis ^7^, the precise mechanisms underlying cardiorenal dysfunction and their interconnection remain largely unknown.

MicroRNAs (miRNAs) are small non-coding RNAs of 17-25 nucleotides that regulate gene expression at a posttranscriptional level by binding to target messenger RNAs or long-coding RNAs ^8–10^. In recent years, miRNA dysregulation has been related to an increasing number of human pathologies, including renal diseases ^11–17^. These molecules are present in all tissues and body fluids and can act as intercellular messengers, which makes them promising candidates as therapeutic targets and diagnostic biomarkers ^18^.

In this work, we identify specific circulating miRNAs as putative biomarkers/therapeutic targets for early CKD diagnosis, disease progression, as well as cardiovascular events and mortality in non-dialysis CKD patients.

## Materials and methods

### Ethics approval

This study was conducted in accordance with the Declaration of Helsinki. The research protocol was reviewed and approved by the institutional Health Ethics Commission and Data Protection Officer of Centro Hospitalar Universitário de São João (CHUSJ), Porto, Portugal (code 130/19 and CES 251.14). All donors and patients involved in the study signed the informed consent form.

### Patient samples

The study population includes two cohorts of non-dialysis CKD patients followed up at the Nephrology Department of CHUSJ. The *discovery cohort* comprises 52 CKD patients randomly selected whose samples were collected between November 2014 and August 2018. The *validation cohort* includes samples from 43 non-dialysis CKD patients who underwent a kidney biopsy in the Nephrology Department of CHUSJ for diagnostic purposes. These samples were collected from November 2019 to December 2022. Patients with acute kidney injury, neoplasia diagnosis, recent infections (<1 week), or known psychiatric disorders were excluded. Plasma samples from 38 healthy blood donors, recruited at the Immunohemotherapy Department of CHUSJ, presenting normal renal function were used as controls. The study size comprehended all eligible patients in the designated period of time.

### Baseline and follow-up evaluation

Demographic data, comorbidity information, as well as CKD and cardiovascular-related parameters were obtained from the patient’s medical records by authorized clinicians. Patients from the *discovery cohort* were prospectively followed up for a median of 70 (57-97) months, to evaluate major renal and cardiovascular outcomes including progression of CKD and end-stage renal disease, renal replacement therapy (RRT) initiation after enrolment, hospitalizations, major adverse cardiovascular events and cardiovascular and all-cause mortality. The *validation cohort* was used to validate the results attained from the *discovery cohort* concerning the miRNA relative expression in comparison with the controls. No clinical data are currently available for this cohort.

### Plasma collection and RNA isolation

Peripheral blood was collected in EDTA-blood collection tubes. The plasma fraction was separated by centrifuging at 800 *g*, for 15 min, at 4°C, transferred into a new tube, and centrifuged at 15001*g*, for 101min at 4°C to remove cell debris before being aliquoted and stored at −80°C.

Total RNA was isolated and purified using miRNeasy serum/plasma kit (Qiagen, Hilden, Germany) following the manufactureŕs instructions. Spike-in controls cel-miR-39-3p (*C. elegans* sequence: 5’-UCACCGGGUGUAAAUCAGCUUG-3’, Invitrogen, Waltham, MA, USA) and cel-miR-54-3p (*C. elegans* sequence: 3’-UACCCGUAAUCUUCAUAAUCCGAG-5’, Invitrogen) were added to each sample and used as external controls.

### Reverse transcription and real-time quantitative polymerase chain reaction (RT-qPCR)

RNA was retrotranscribed into cDNA using the TaqMan Advanced miRNA cDNA synthesis kit (Applied Biosystems, Carlsbad, CA, USA) according to the manufacturer’s protocol.

The expression levels of miR-29b-3p, miR-30c-5p, miR-132-3p, miR-146a-5p, miR-199a-5p, and miR-324-3p were analyzed using TaqMan Advanced miRNA assays (Applied Biosystems). RT-qPCR was carried out in 7500 Fast Real-Time PCR System (Applied Biosystems) using cDNA, TaqMan Fast Advanced Master Mix (Applied Biosystems), and TaqMan Advanced microRNA Assays specific for each miRNA (Applied Biosystems). Each reaction was performed in duplicates, and samples with replicate differences higher than 0.5 were not considered for further analysis. Relative miRNA expression was calculated using the ΔCq method according to the Minimum Information for Publication of Quantitative Real-Time PCR Experiments (MIQE) guidelines ^19^.

### Statistical analysis

Statistical analysis was performed using SPSS 26 (IBM, Manhattan, NY, USA) and GraphPad Prism 9.0.1 (GraphPad Software, San Diego, CA, USA) statistical software. Shapiro-Wilk test was performed to study the distribution of the data. Mann-Whitney and Kruskal-Wallis tests were carried out to assess differences in continuous variables between two or more groups, respectively. Differences between categorical variables were evaluated using Fisheŕs exact test. Correlations between numerical variables were analyzed by using Spearmańs correlation coefficient. Receiver operating characteristics (ROC) curves were performed to assess the discriminatory diagnostic power of a variable. The specificity and sensitivity percentages displayed correspond to the maximum Youden index (calculated as sensitivity + specificity - 1). Significant differences between AUC values were assessed with R Studio 2023.12.1+402 (R studio software, PBC, Boston, USA). Logistic binary regression was performed to predict the occurrence of a categorical variable based on predictor variables. In all cases, statistical significance was considered when *P*-value < 0.05. Whenever there was missing data, the particular patients were excluded from the analysis.

## Results

### Study population

Baseline demographic data, comorbidity information, as well as CKD and cardiovascular-related parameters are summarized in Table 1. Demographic data shows that CKD patients were significantly older than the control group, and as expected, patients from both CKD groups exhibited higher creatinine levels when compared to the control group. No significant differences were found regarding gender or CKD etiology. The most predominant CKD etiology in both the *discovery* and the *validation cohorts* was IgA nephropathy (IgAN) (20.41% and 34.49%, respectively) (Figure 1).

**Figure 1.**
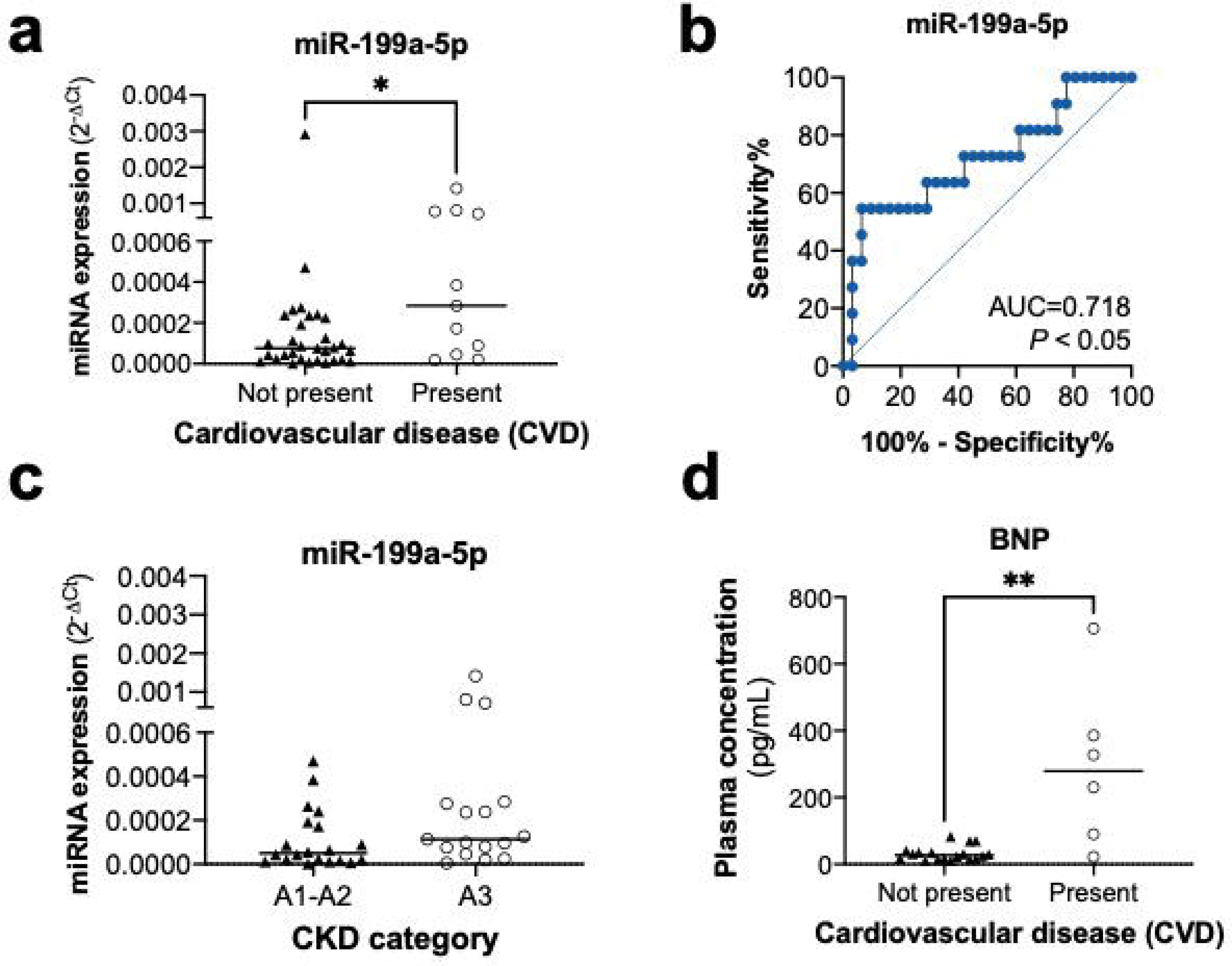
Etiology of CKD patients. Graphs show the percentage of CKD causes in the (a) *discovery* and (b) *validation cohorts*. CKD: chronic kidney disease.

**Table 1.**
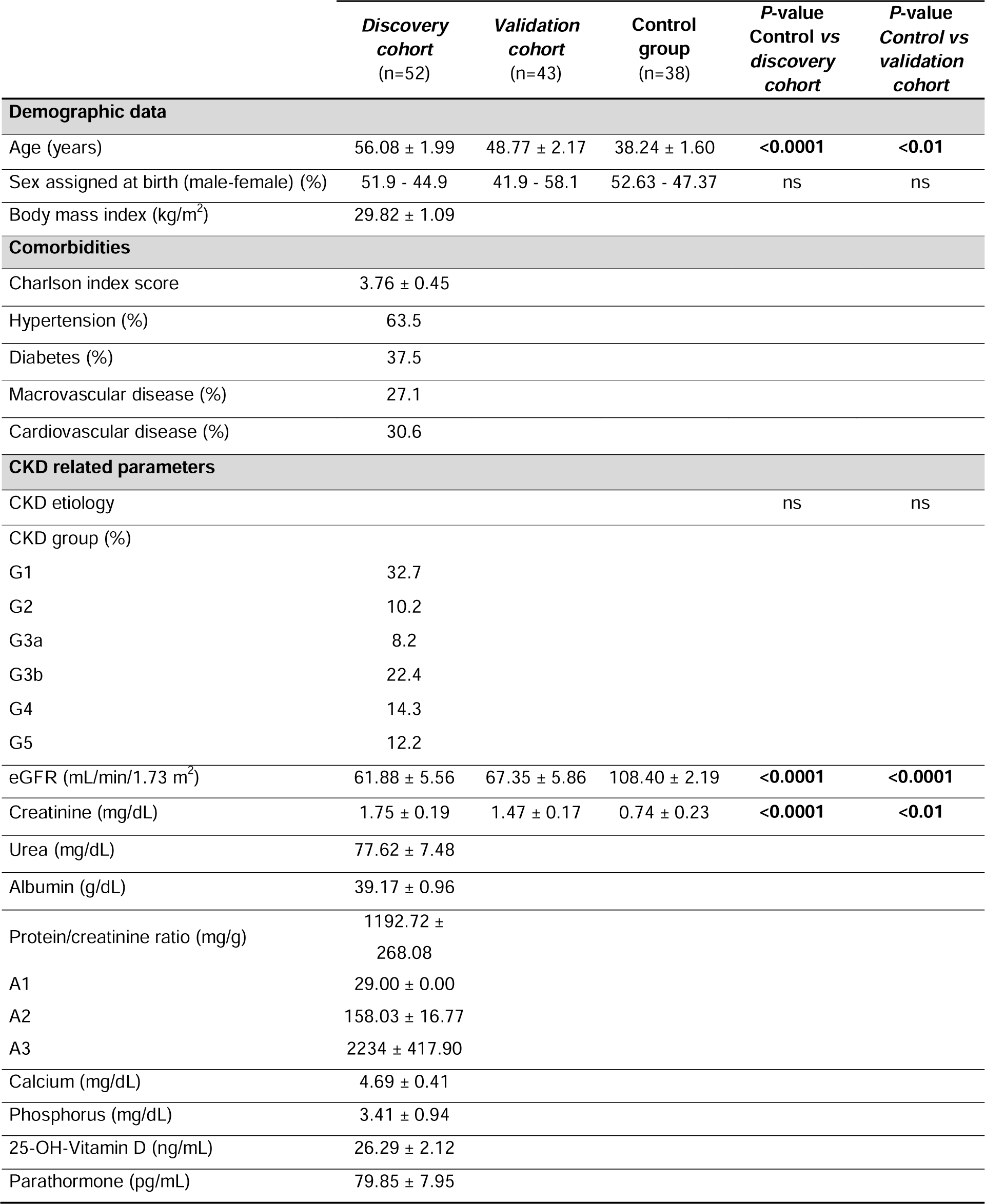
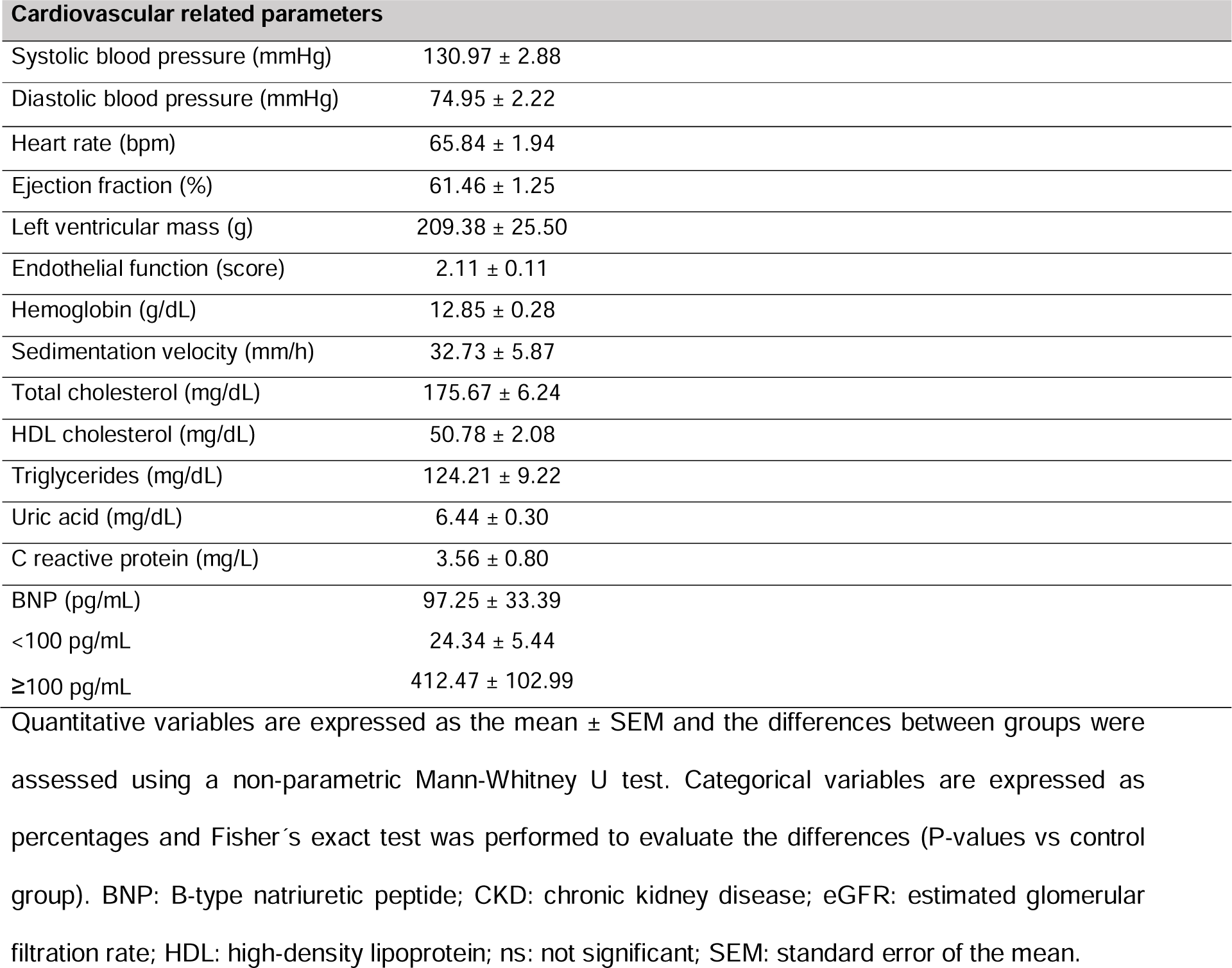
Baseline demographic information and clinical data.

|  | *Discovery cohort*<br>(n=52) | *Validation cohort*<br>(n=43) | *Control group*<br>(n=38) | *P-value*<br>*Control vs*<br>*discovery cohort* | *P-value*<br>*Control vs*<br>*validation cohort* |
| --- | --- | --- | --- | --- | --- |
| **Demographic data** |  |  |  |  |  |
| Age (years) | 56.08 ± 1.99 | 48.77 ± 2.17 | 38.24 ± 1.60 | <0.0001 | <0.01 |
| Sex assigned at birth (male-female) (%) | 51.9 - 44.9 | 41.9 - 58.1 | 52.63 - 47.37 | ns | ns |
| Body mass index (kg/m<sup>2</sup>) | 29.82 ± 1.09 |  |  |  |  |
| **Comorbidities** |  |  |  |  |  |
| Charlson index score | 3.76 ± 0.45 |  |  |  |  |
| Hypertension (%) | 63.5 |  |  |  |  |
| Diabetes (%) | 37.5 |  |  |  |  |
| Macrovascular disease (%) | 27.1 |  |  |  |  |
| Cardiovascular disease (%) | 30.6 |  |  |  |  |
| **CKD related parameters** |  |  |  |  |  |
| CKD etiology |  |  |  | ns | ns |
| CKD group (%) |  |  |  |  |  |
| G1 | 32.7 |  |  |  |  |
| G2 | 10.2 |  |  |  |  |
| G3a | 8.2 |  |  |  |  |
| G3b | 22.4 |  |  |  |  |
| G4 | 14.3 |  |  |  |  |
| G5 | 12.2 |  |  |  |  |
| eGFR (mL/min/1.73 m<sup>2</sup>) | 61.88 ± 5.56 | 67.35 ± 5.86 | 108.40 ± 2.19 | <0.0001 | <0.0001 |
| Creatinine (mg/dL) | 1.75 ± 0.19 | 1.47 ± 0.17 | 0.74 ± 0.23 | <0.0001 | <0.01 |
| Urea (mg/dL) | 77.62 ± 7.48 |  |  |  |  |
| Albumin (g/dL) | 39.17 ± 0.96 |  |  |  |  |
| Protein/creatinine ratio (mg/g) | 1192.72 ± 268.08 |  |  |  |  |
| A1 | 29.00 ± 0.00 |  |  |  |  |
| A2 | 158.03 ± 16.77 |  |  |  |  |
| A3 | 2234 ± 417.90 |  |  |  |  |
| Calcium (mg/dL) | 4.69 ± 0.41 |  |  |  |  |
| Phosphorus (mg/dL) | 3.41 ± 0.94 |  |  |  |  |
| 25-OH-Vitamin D (ng/mL) | 26.29 ± 2.12 |  |  |  |  |
| Parathormone (pg/mL) | 79.85 ± 7.95 |  |  |  |  |

**Cardiovascular related parameters**
|  |  |
| --- | --- |
| Systolic blood pressure (mmHg) | 130.97 ± 2.88 |
| Diastolic blood pressure (mmHg) | 74.95 ± 2.22 |
| Heart rate (bpm) | 65.84 ± 1.94 |
| Ejection fraction (%) | 61.46 ± 1.25 |
| Left ventricular mass (g) | 209.38 ± 25.50 |
| Endothelial function (score) | 2.11 ± 0.11 |
| Hemoglobin (g/dL) | 12.85 ± 0.28 |
| Sedimentation velocity (mm/h) | 32.73 ± 5.87 |
| Total cholesterol (mg/dL) | 175.67 ± 6.24 |
| HDL cholesterol (mg/dL) | 50.78 ± 2.08 |
| Triglycerides (mg/dL) | 124.21 ± 9.22 |
| Uric acid (mg/dL) | 6.44 ± 0.30 |
| C reactive protein (mg/L) | 3.56 ± 0.80 |
| BNP (pg/mL) | 97.25 ± 33.39 |
| <100 pg/mL | 24.34 ± 5.44 |
| ≥100 pg/mL | 412.47 ± 102.99 |

Quantitative variables are expressed as the mean ± SEM and the differences between groups were assessed using a non-parametric Mann-Whitney U test. Categorical variables are expressed as percentages and Fisher's exact test was performed to evaluate the differences (P-values vs control group). BNP: B-type natriuretic peptide; CKD: chronic kidney disease; eGFR: estimated glomerular filtration rate; HDL: high-density lipoprotein; ns: not significant; SEM: standard error of the mean.

### Circulating miR-30c-5p and miR-132-3p are downregulated in plasma samples from CKD patients

From the panel of 6 miRNAs investigated, miR-30c-5p and miR-132-3p were found to be significantly decreased in both the *discovery* and the *validation cohorts,* in comparison with healthy controls (*P* < 0.0001 and *P* < 0.05, respectively; Figure 2). The results for miR-29b-3p, miR-146a-5p, miR-199a-5p, and miR-324-3p expression in comparison with the control group were not validated in the *validation cohort* and were excluded from follow-up analysis (Figure S1). The analysis of putative correlations between miRNA expression and the patient’s demographic data showed a correlation between miR-132-3p and age in the *discovery cohort.* However, no significant correlations were identified concerning the body mass index (BMI) (Table 2). Considering that age was a significant factor impacting the levels of miR-132-3p expression in the *discovery cohort*, a regression analysis was performed to adjust the significance for the variable age. The adjusted analysis reaffirmed the former identified differences between miR-132-5p expression in CKD *vs* controls (Table 3). Finally, an evaluation of data concerning the causes of CKD was also performed, and the analysis revealed no significant differences in miR-30c-5p or miR-132-3p expression regarding disease etiology in both the *discovery* and the *validation cohorts* (Table 4).

**Figure 2.**
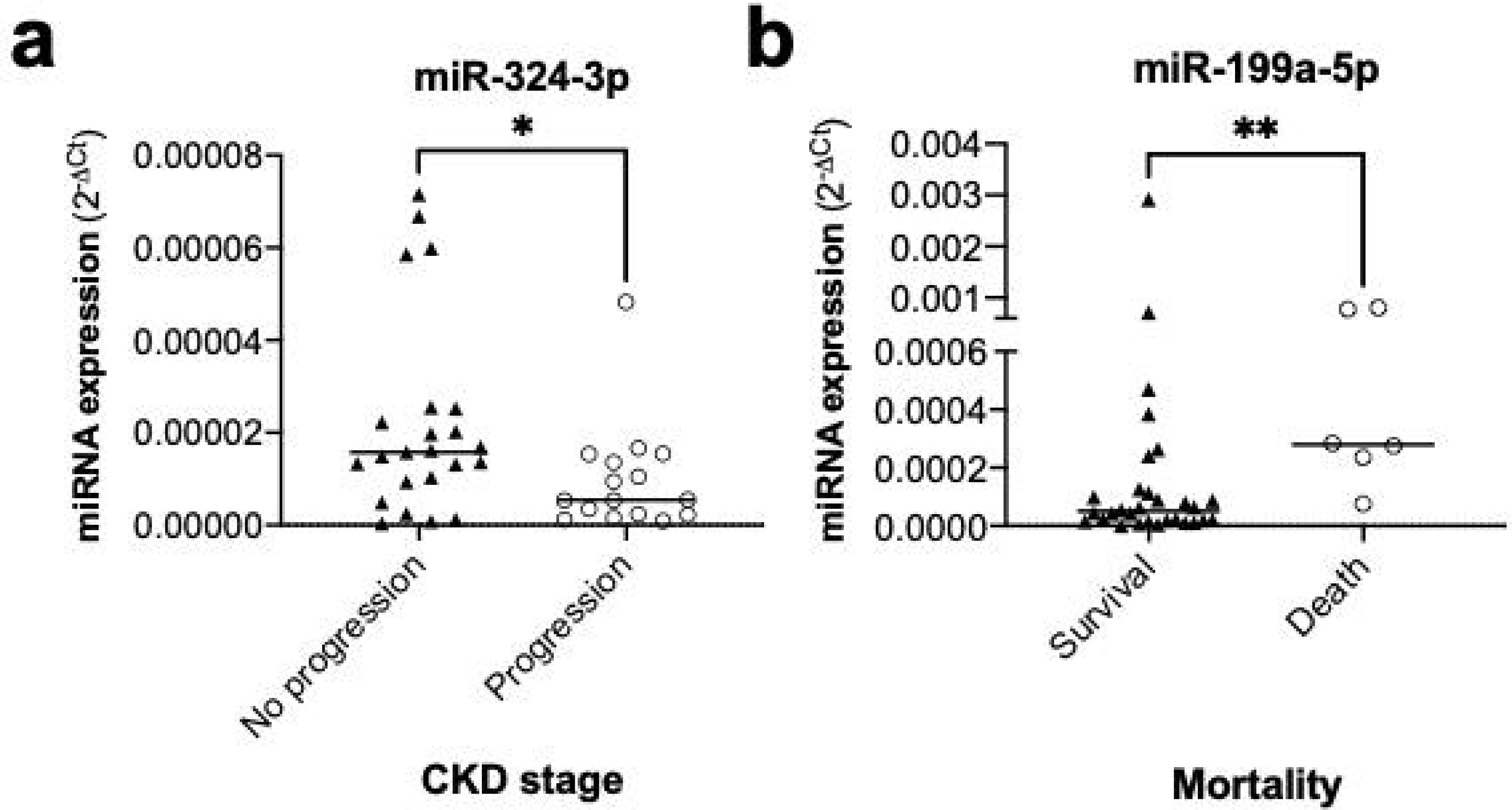
Circulating miR-30c-5p and miR-132-3p are downregulated in plasma samples from CKD patients from both (a) the d*iscovery* (n=52) and (b) the *validation* cohorts (n=43), in comparison with the control group (n=38). miRNA quantification was assessed by real-time qRT-PCR (median; Mann-Whitney test; \**P* < 0.05 and \*\*\*\**P* < 0.0001 *vs* control group). CKD: chronic kidney disease; miRNA: microRNA.

**Table 2.** Correlation between miR-30c-5p and miR-132-3p expression with age and body mass index.

| miRNA |  | Discovery cohort<br>(n=52) |  | Validation cohort<br>(n=43) |
| --- | --- | --- | --- | --- |
|  |  | Age | BMI | Age |
| miR-30c-5p | $\rho$ | -0.102 | 0.037 | -0.191 |
|  | *P*-value | 0.491 | 0.836 | 0.108 |
| miR-132-3p | $\rho$ | -0.322 | 0.126 | -0.041 |
|  | *P*-value | **0.029\*** | 0.485 | 0.724 |

Spearman's Rho coefficient ( $\rho$ ) and corresponding *P*-values for each miRNA are shown as the result of the correlation analysis (\* $P < 0.05$ ). BMI: body mass index; CKD: chronic kidney disease; miRNA: microRNA.

**Table 3.** Statistical differences for miR-132-5p expression between the control group (n=38) and CKD patients from the *discovery cohort* (n=52) with and without adjusting to age.

| miRNA | *P*-value<br>(without age adjustment) | *P*-value<br>(adjusted to age) |
| --- | --- | --- |
| miR-132-3p | 0.000* | **0.016\*** |

\* $P < 0.05$ , CKD: chronic kidney disease; miRNA: microRNA.

**Table 4.** Differences in miR-30c-5p and miR-132-3p expression regarding the etiology of the disease in plasma samples from CKD patients.

| miRNAs | Discovery cohort<br>(n=49) | Validation cohort<br>(n=39) |
| --- | --- | --- |
| miR-30c-5p | 0.556 | 0.442 |
| miR-132-3p | 0.208 | 0.324 |

*P*-values resulting from performing a Mann-Whitney test are shown for each miRNA. CKD: chronic kidney disease; miRNAs: microRNAs.

### Diagnostic value of miR-30c-5p and miR-132-3p for CKD

To assess the diagnostic value of the differentially expressed miRNAs for CKD, ROC curves were performed using the data obtained from the *discovery* cohort independently of the disease stage. As displayed in Figure 3, the ROC curve analysis of miR-30c-5p showed an area under the curve (AUC) of 0.832 (*P* < 0.001, 95% CI: 0.743-0.921), with a maximum value of Youden index of 0.594, indicating the cut-off points that determine the highest sensitivity and specificity (76.5% and 82.9% respectively). MiR-132-3p presents a 91.8% of sensitivity and 78.9% of specificity for a Youden index of 0.707, with an AUC of 0.856 (*P* < 0.001, 95% CI 0.766-0.945). When combined, miR-30c-5p and miR-132-3p exhibit an AUC value of 0.864 (*P* < 0.001, 95% CI 0.776-0.952), demonstrating no significant improvement compared to individual miRNAs. In comparison to the diagnostic value of eGFR (AUC=0.823, *P* < 0.0001), the combination of eGFR with miR-30c-5p (AUC=0.901, *P* < 0.0001), miR-132-3p (AUC=0.897, *P* < 0.0001) and the combination of the expression of both miRNAs (AUC=0.917, P < 0.0001) exhibited a significantly higher AUC value (*P* < 0.05, *P* < 0.01 and *P* < 0.05, respectively), that serves as a quantitative measure of accuracy in discriminating CKD.

**Figure 3.**
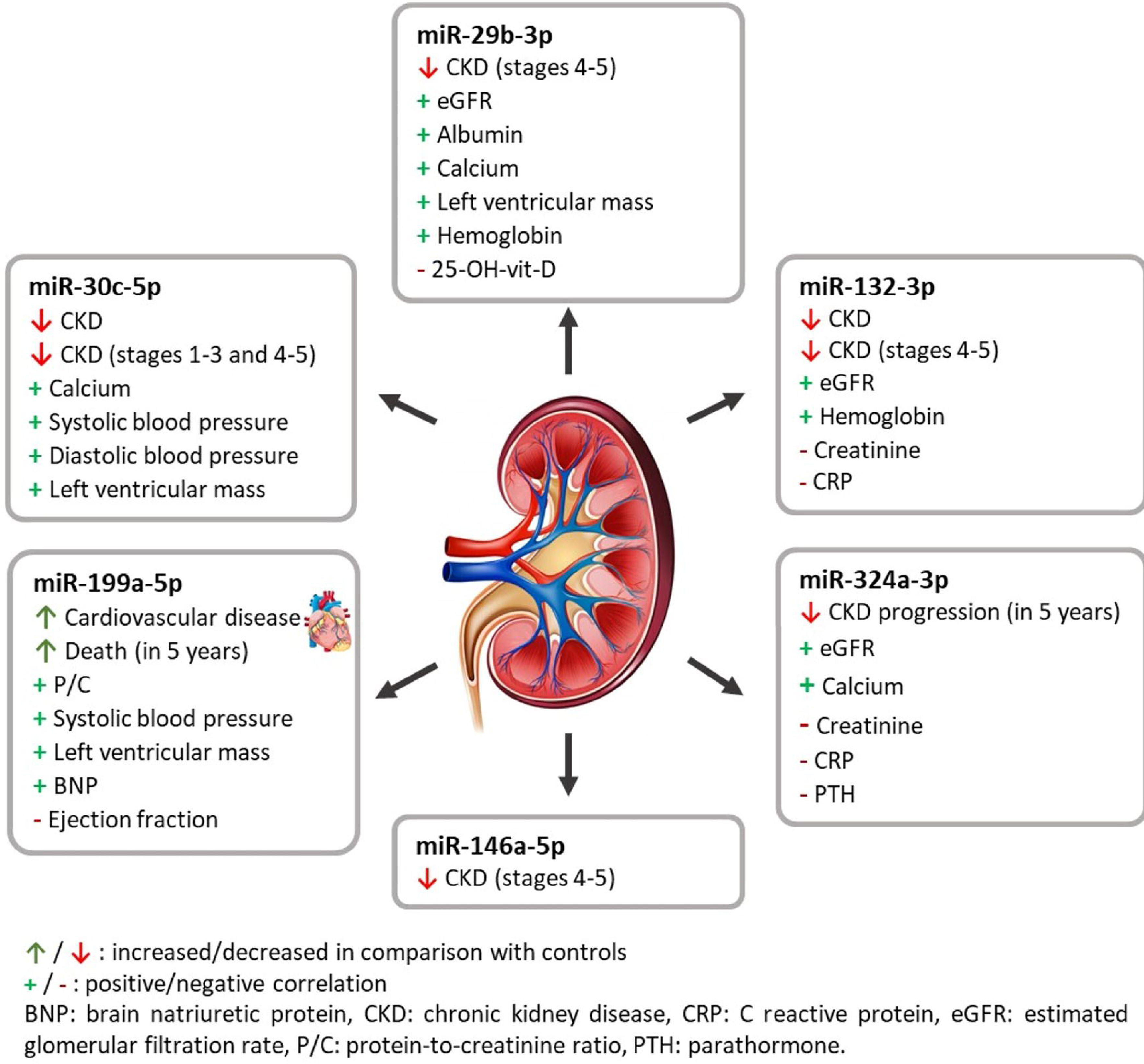
Specificity and sensitivity of miR-30c-5p, miR-132-3p, eGFR and their combination for CKD diagnosis. ROC curves concerning (a) miR-30c-5p, (b) miR-132-3p, (c) their combined expression, as well as (d) eGFR, and the combined diagnostic ability of eGFR with (e) miR-30c-5p, (f) miR-132-3p, and with (g) miR-30c-5p plus miR-132-5p were determined. AUC: area under the curve; CKD: chronic kidney disease; eGFR: estimated glomerular filtration rate; miRNA: microRNA; ROC: Receiver operation characteristic.

### Circulating miR-30c-5p as a biomarker for early CKD diagnosis

Considering the potential clinical relevance of identifying a biomarker that allows the diagnosis of CKD during the initial stages, we analyzed the expression of circulating miRNAs in early CKD (G1-G2 and G1-G3). Results showed that miR-30c-5p levels were downregulated in plasma samples from CKD patients in stages G1-G3 in both the *discovery* and the *validation* cohort compared to healthy controls (Figure 4a and 4b). The restriction of the analysis to G1-G2 displayed the same tendency of decrease in G1-G3 in comparison with the control group, but miR-30c was only significantly downregulated in the *discovery* cohort (Figure 4f) and results have not reached statistical significance in the *validation* cohort (Figure 4g). ROC curve analysis of miR-30c-5p expression showed an AUC of 0.807, with 71.4% sensitivity and 82.9% specificity in discriminating between the control group and CKD G1-G3 (*P* < 0.001, 95% CI: 0.704-0.911) for a Youden index of 0.543.

**Figure 4.**
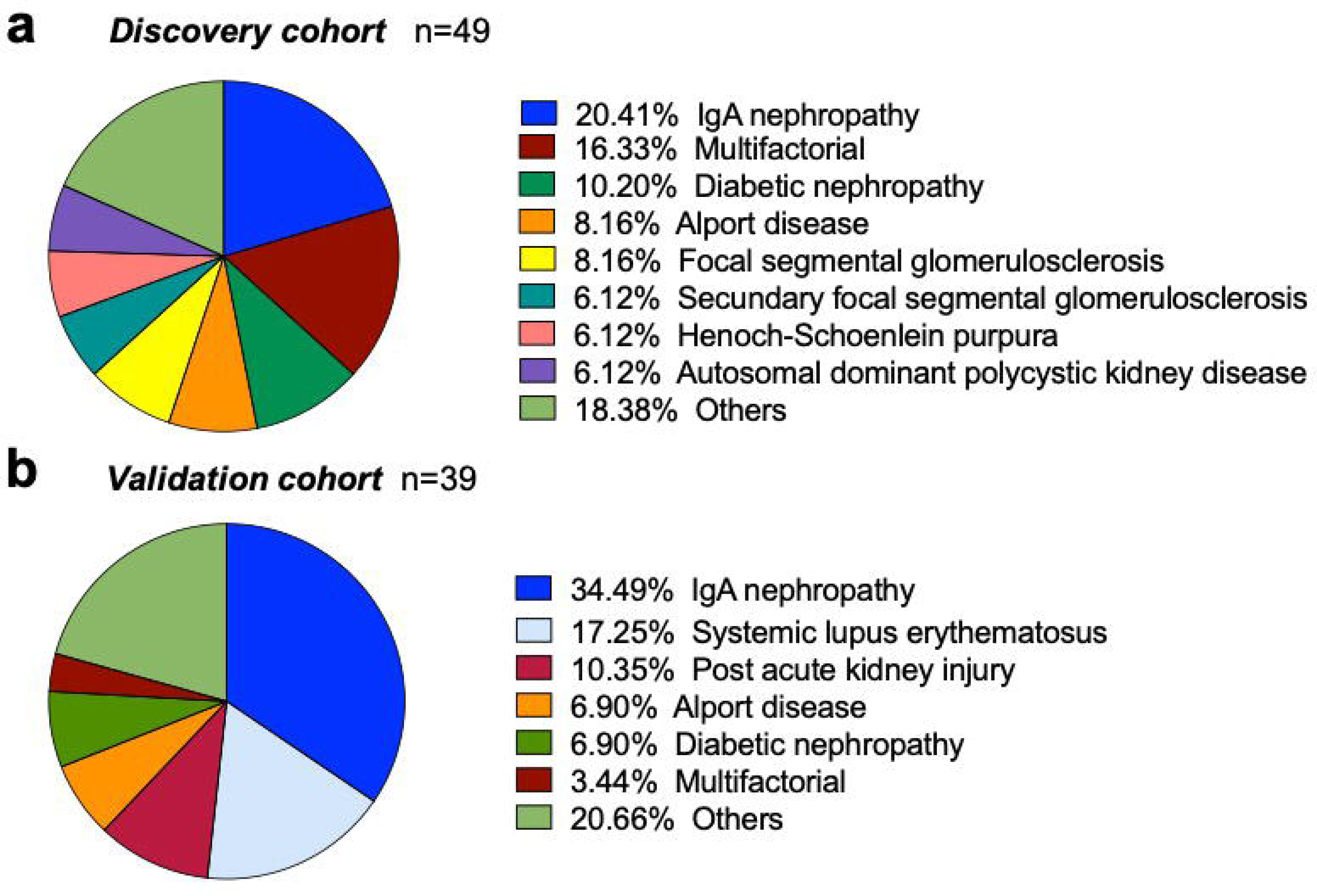
Circulating miR-30c-5p is downregulated in plasma samples from early CKD patients (G1-G3) in both (a) the *discovery* (n=35) and (b) the *validation* (n=34) cohorts compared with the control group (n=38). miRNA expression was determined by real-time qRT-PCR (median; Mann-Whitney test; \**P* < 0.05 and \*\*\*\**P* < 0.0001 *vs* control group). ROC curves show the diagnostic ability to discriminate CKD at the early stages of the disease of (c) miR-30c-5p expression, (d) eGFR, and (e) the combination of both parameters measured in the *discovery cohort*. The restriction of the analysis to CKD G1-G2 portraited a significant downregulation of miR-30c-5p only in the *discovery cohort (f)* (n=35) that was not statistically significant in the *validation cohort* (g) (n=34), compared with the control group (n=38). AUC: area under the curve; CKD: chronic kidney disease; eGFR: estimated glomerular filtration rate; miRNA: microRNA; ROC: Receiver operation characteristic.

Furthermore, the AUC of the combination of eGFR data with miRNA-30c-5p expression (AUC=0.866, *P* < 0.0001) was found to be significantly higher (*P* < 0.05) than the AUC obtained from the eGFR ROC curve (AUC=0.759, *P* < 0.0001) (Figure 4c, 4d and 4e). In addition, logistic regression analysis showed that miR-30c-5p was significant in predicting CKD in the early stages (G1-G3), correctly classifying 85.7% of the pathological cases. The odd ratio (OR) value suggests that miR-30c-5p exerts a protective effect as higher levels would be associated with a decreased probability of CKD development (Table 5).

**Table 5.**
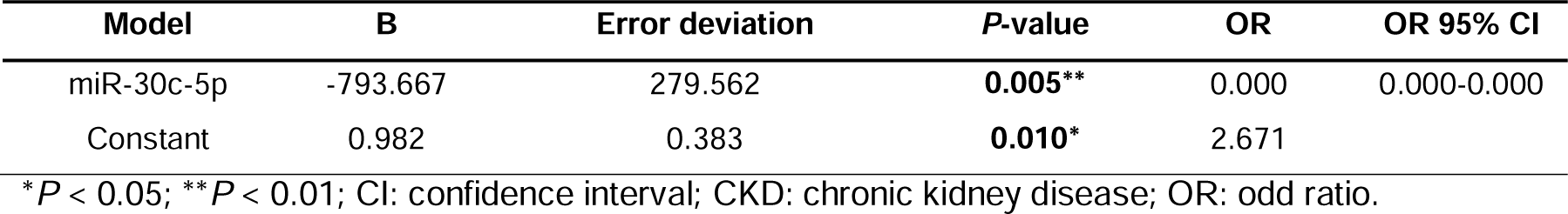
Logistic binary regression model to predict early CKD based on the expression of miR-30c-5p in plasma samples from the *discovery cohort*.

| Model | B | Error deviation | *P*-value | OR | OR 95% CI |
| --- | --- | --- | --- | --- | --- |
| miR-30c-5p | -793.667 | 279.562 | **0.005\*\*** | 0.000 | 0.000-0.000 |
| Constant | 0.982 | 0.383 | **0.010\*** | 2.671 |  |

\* $P < 0.05$ ; \*\* $P < 0.01$ ; CI: confidence interval; CKD: chronic kidney disease; OR: odd ratio.

### Circulating miR-30c-5p, miR-132-3p, miR-29b-3p, and miR-146a-5p are downregulated in advanced CKD stages

Next, we evaluated circulating miRNA expression in CKD patients with more advanced stages of the disease (G4 and G5). Results showed a significant decrease in miR-30c-5p, miR-132-3p, miR-29b-3p, and miR-146-5p levels in plasma samples from CKD patients from both the *discovery* and the *validation* cohorts when compared to the control group (healthy donors) (Figure 5).

**Figure 5.**
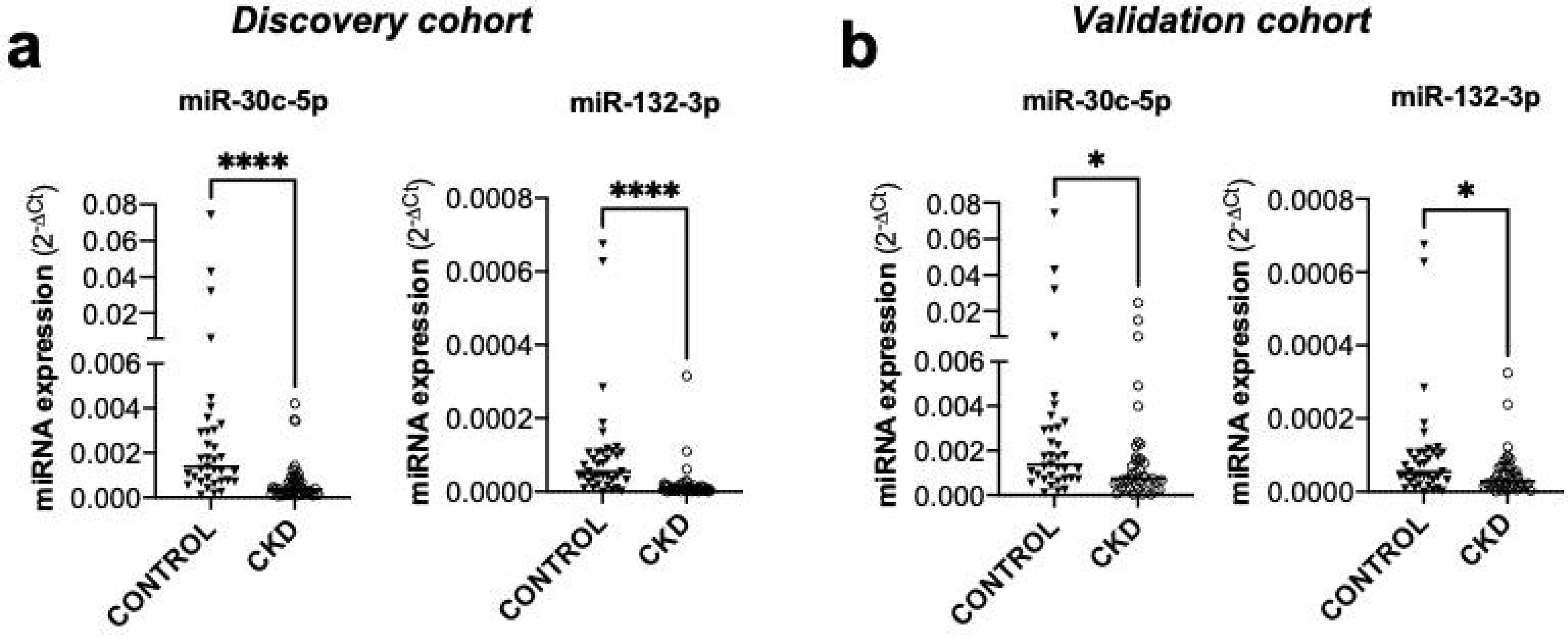
Circulating miR-30c-5p, miR-132-3p, miR-29b-3p, and miR-146a-5p are downregulated in plasma samples from patients with advanced CKD (G4-G5) compared with the control group. Graphs show miRNA expression assessed in plasma from 38 controls and CKD patients in stages G4 to G5 from both (a) the *discovery* (n=13) and (b) the *validation* (n=6) cohorts by real-time qRT-PCR (median; Mann-Whitney test; \**P* < 0.05, \*\**P* < 0.01, \*\*\*\**P* < 0.0001 *vs* control group). AUC: area under the curve; CKD: chronic kidney disease; miRNA: microRNA; ROC: Receiver operation characteristic.

### Correlation between circulating microRNAs and clinical parameters in CKD

The implication of circulating miRNAs in renal function was investigated concerning putative correlations with the clinical data. These studies were conducted exclusively in data from the *discovery cohort* due to the unavailability of clinical data from the *validation cohort*. As shown in Table 6, eGFR correlates positively with miR-132-3p, miR-29-3p, and miR-324-3p. Moreover, a positive correlation was observed between serum albumin content and miR-29-3p. Concerning protein-to-creatinine ratio (P/C), a positive correlation was identified with miR-199a-5p.

**Table 6.** Correlation between circulating miRNAs and clinical data of CKD patients.

| miRNAs |  | miR-30c-5p | miR-132-3p | miR-29b-3p | miR-146a-5p | miR-199a-5p | miR-324-3p |
| --- | --- | --- | --- | --- | --- | --- | --- |
| **Renal function** |  |  |  |  |  |  |  |
| **eGFR**<br>(mL/min/1.73 m<sup>2</sup>) | $\rho$ | 0.061 | 0.381 | 0.300 | 0.256 | -0.109 | 0.407 |
|  | *P*-value | 0.678 | 0.009** | 0.036* | 0.079 | 0.492 | 0.006** |
| **Creatinine**<br>(mg/dL) | $\rho$ | 0.053 | -0.306 | -0.167 | -0.209 | 0.153 | -0.356 |
|  | *P*-value | 0.718 | 0.039* | 0.250 | 0.154 | 0.332 | 0.016* |
| **Urea** (mg/dL) | $\rho$ | 0.022 | -0.274 | -0.229 | -0.207 | 0.188 | -0.290 |
|  | *P*-value | 0.885 | 0.068 | 0.118 | 0.162 | 0.239 | 0.056 |
| **Albumin** (g/dL) | $\rho$ | 0.272 | 0.258 | 0.359 | 0.114 | -0.097 | 0.185 |
|  | *P*-value | 0.061 | 0.084 | 0.011* | 0.439 | 0.539 | 0.225 |
| **P/C** (mg/g) | $\rho$ | 0.173 | 0.063 | -0.050 | 0.118 | 0.368 | -0.167 |
|  | *P*-value | 0.306 | 0.716 | 0.767 | 0.488 | 0.035* | 0.330 |
| **Inflammation** |  |  |  |  |  |  |  |
| **CRP** (mg/L) | $\rho$ | -0.040 | -0.532 | -0.269 | -0.090 | -0.001 | -0.479 |
|  | *P*-value | 0.845 | 0.005** | 0.176 | 0.662 | 0.995 | 0.015* |
| **Sedimentation velocity** (mm/h) | $\rho$ | 0.210 | -0.240 | 0.005 | -0.125 | 0.080 | -0.217 |
|  | *P*-value | 0.403 | 0.322 | 0.983 | 0.620 | 0.753 | 0.386 |
| **Mineral metabolism** |  |  |  |  |  |  |  |
| **25-OH-vit-D**<br>(ng/mL) | $\rho$ | -0.239 | 0.012 | -0.355 | -0.270 | -0.216 | -0.240 |
|  | *P*-value | 0.180 | 0.947 | 0.040* | 0.128 | 0.260 | 0.186 |
| **Calcium** (mg/dL) | $\rho$ | 0.309 | 0.225 | 0.386 | 0.231 | 0.029 | 0.348 |
|  | *P*-value | 0.049* | 0.163 | 0.012* | 0.145 | 0.869 | 0.028* |
| **PTH** (pg/mL) | $\rho$ | -0.054 | -0.228 | -0.178 | -0.151 | 0.063 | -0.422 |
|  | *P*-value | 0.748 | 0.169 | 0.278 | 0.365 | 0.724 | 0.009** |
| **Phosphorus**<br>(mg/dL) | $\rho$ | 0.138 | -0.171 | -0.052 | 0.019 | 0.122 | -0.160 |
|  | *P*-value | 0.389 | 0.292 | 0.744 | 0.904 | 0.480 | 0.324 |

Spearman's rho coefficient ( $\rho$ ) and corresponding *P*-values for each miRNA are shown as the result of the correlation analysis (\**P* < 0.05; \*\**P* < 0.01). CKD: chronic kidney disease; eGFR: estimated glomerular filtration rate; miRNAs: microRNAs; P/C: protein/creatinine ratio; PTH: parathormone; vit- D: vitamin D.

Furthermore, considering this parameter to discriminate patients in albuminuria categories A1-A2 and A3 (considering P/C ratio and the threshold bellow and above 500mg/g, respectively) a tendency of increase of miR-199a-5p in the worse category (*P* = 0.0603; Figure 6 c) was identified. No differences regarding such differentiation analysis of the P/C ratio were verified for any of the other miRNAs (data not shown).

**Figure 6.**
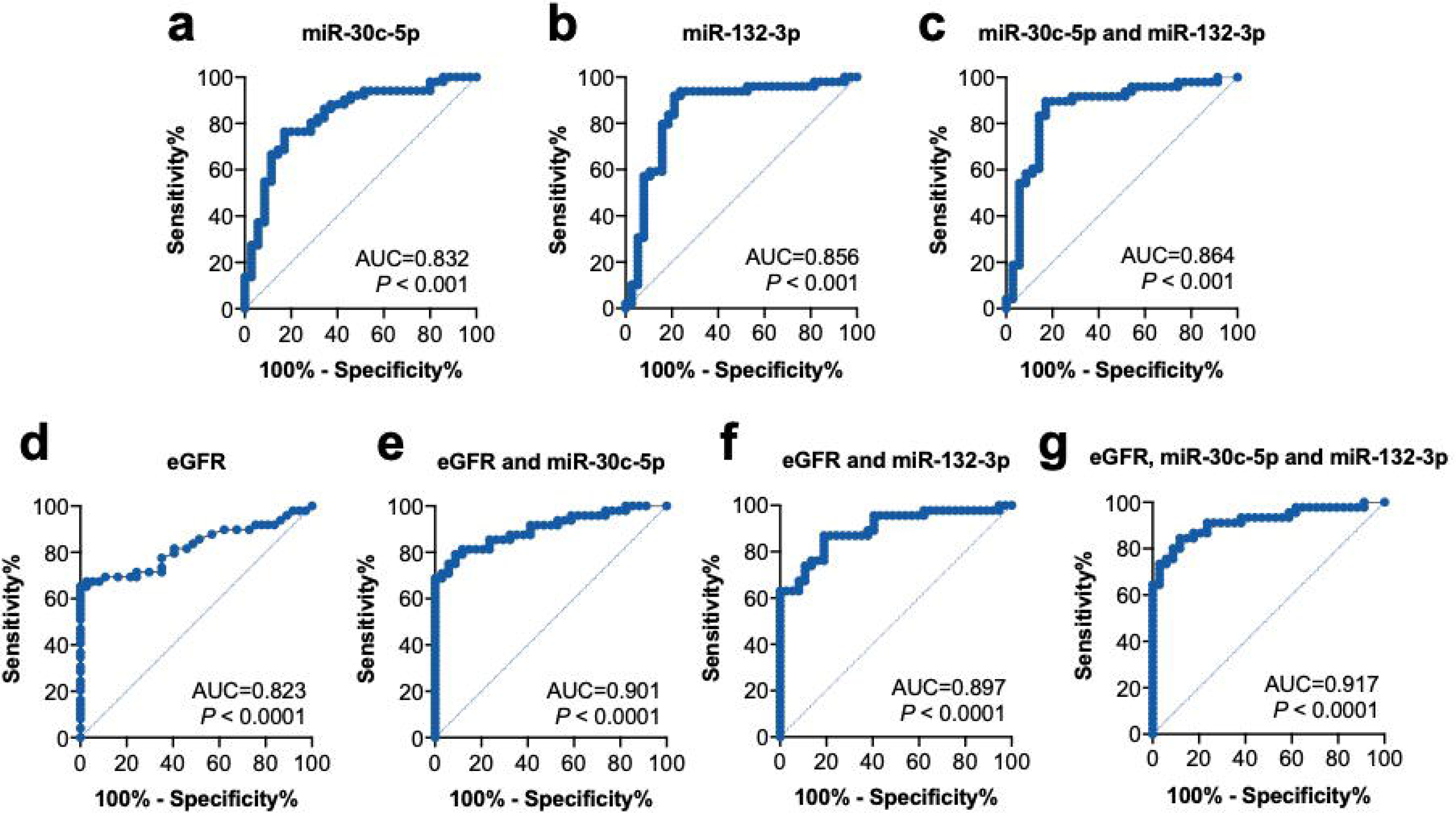
(a) Circulating miR-199a-5p is increased in CKD patients who suffer from cardiovascular disease (n=11) in comparison with the ones that do not (n=31). miRNAs were quantified by real-time qRT-PCR (median; Mann Whitney test; \**P* < 0.05). (b) ROC analysis of miR-199a-5p shows a significative diagnostic power to discriminate CKD patients without and with cardiovascular diseases. (c) The segmentation of patients according to P/C ratio shows a tendency of increased expression of miR-199a-5p in patients with P/C > 500 mg/mL (n=20, CKD categories A1-A2) in comparison with patients with P/C < 500 mg/mL; (n=17, CKD category A3) (median, Mann Whitney test, *P* = 0.0603). (d) BNP quantification discriminated by the absence or presence of cardiovascular disease shows a significant increase in the last group (median; Mann Whitney test; \**P* < 0.05, \*\**P* < 0.01). AUC: area under the curve; BNP: brain natriuretic peptide; CKD: chronic kidney disease; miRNA: microRNA; P/C: Urine protein/creatinine ratio; ROC: Receiver operation characteristic.

Progression of CKD is usually characterized by an increase in the inflammatory state and a dysregulation of mineral metabolism. Hence, we also explored the potential relationship between miRNA expression and systemic inflammation. Our results disclosed a negative correlation between plasma miR-132-3p and miR-324-3p with blood levels of the inflammatory marker C reactive protein (CRP). No significant correlations were found between miRNA levels and sedimentation velocity.

Regarding mineral metabolic alterations, we observed a negative correlation between miR-29-3p and miR-324-3p levels with vitamin D and the secretion of parathormone, respectively. Moreover, a positive correlation was found between calcium blood levels and miR-30c-5p, miR-29b-3p, and miR-324-3p expression.

### Circulating miR-199a-5p is associated with CVD in CKD

The main risk factors for developing CKD are diabetes and hypertension, followed by cardiovascular diseases and a family history of kidney failure. Based on this information, we evaluated differences in miRNA expression regarding the most common comorbidities of CKD. As shown in Table 7, miR-199a-5p was associated with cardiovascular disease in CKD patients. No significant differences were observed in miRNA levels regarding the remaining comorbidities. In particular, miR-199a-5p was upregulated in CKD patients with CVD (Figure 6a) and ROC curve analysis showed an AUC of 0.7185 (*P* < 0.05, 95% CI: 0.528-0.909) with a 54.5% sensitivity and a 93.5% specificity (for a Youden index of 0.48) for discriminating these patients (Figure 6b).

**Table 7.** Differences in miRNA expression regarding the main comorbidities associated with CKD.

| Comorbidity | Diabetes<br>(n=48) | Hypertension<br>(n=48) | Cardiovascular<br>Disease<br>(n=49) | Macrovascular<br>disease<br>(n=48) |
| --- | --- | --- | --- | --- |
| miR-30c-5p | 0.393 | 0.963 | 0.764 | 0.831 |
| miR-132-3p | 1.000 | 0.485 | 0.360 | 0.093 |
| miR-29-3p | 0.639 | 0.085 | 0.298 | 0.167 |
| miR-146a-5p | 0.965 | 0.693 | 0.868 | 0.849 |
| miR-199a-5p | 0.055 | 0.072 | 0.033* | 0.230 |
| miR-324-3p | 0.172 | 0.279 | 0.395 | 0.169 |

*P*-value obtained after performing a Mann-Whitney test (\**P* < 0.05). CKD: chronic kidney disease;
miRNA: microRNA.

Concerning clinical data associated with cardiovascular function (Table 8), a positive correlation between miR-30c-5p and both systolic and diastolic blood pressure levels was observed, while miR-199a-5p exhibited a positive correlation specifically with systolic blood pressure. miR-199a-5p was also negatively correlated with ejection fraction and positively correlated with left ventricular mass, which contrasted with the positive correlation identified for miR-30c-5p and miR-29-3p. Another common consequence of CKD is the development of anemia, especially among patients in the advanced stages of the disease. In relation, hemoglobin levels were positively correlated with miR132-3p and miR-29-3p expression. Lastly, a positive correlation between miR-199a-5p expression and brain natriuretic peptide (BNP) was also identified and BNP levels were found to be increased in CKD patients with CVD (Figure 6d).

**Table 8.** Correlation between plasmatic miRNA expression and clinical parameters related to the cardiovascular system.

| miRNAs |  | miR-30c-5p | miR-132-3p | miR-29-3p | miR-146a-5p | miR-199a-5p | miR-324-3p |
| --- | --- | --- | --- | --- | --- | --- | --- |
| **Systolic blood pressure**<br>(n=41) | $\rho$ | 0.396 | 0.138 | 0.138 | 0.089 | 0.397 | 0.110 |
|  | *P*-value | 0.011* | 0.410 | 0.390 | 0.585 | 0.018* | 0.507 |
| **Diastolic blood pressure**<br>(n=41) | $\rho$ | 0.355 | 0.270 | 0.278 | 0.171 | 0.282 | 0.138 |
|  | *P*-value | 0.025* | 0.102 | 0.078 | 0.293 | 0.101 | 0.402 |
| **Heart rate**<br>(n=38) | $\rho$ | -0.312 | -0.248 | -0.145 | -0.217 | -0.211 | -0.083 |
|  | *P*-value | 0.057 | 0.150 | 0.384 | 0.191 | 0.238 | 0.628 |
| **Ejection fraction**<br>(n=13) | $\rho$ | -0.106 | -0.274 | -0.357 | -0.254 | -0.640 | -0.476 |
|  | *P*-value | 0.743 | 0.365 | 0.231 | 0.425 | 0.046* | 0.100 |
| **Left ventricular mass**<br>(n=13) | $\rho$ | 0.595 | 0.465 | 0.575 | 0.455 | 0.770 | -0.030 |
|  | *P*-value | 0.041* | 0.109 | 0.040* | 0.137 | 0.009** | 0.922 |
| **Endothelial function**<br>(n=43) | $\rho$ | -0.179 | 0.007 | 0.003 | -0.252 | -0.234 | -0.002 |
|  | *P*-value | 0.258 | 0.967 | 0.985 | 0.107 | 0.163 | 0.990 |
| **Hemoglobin**<br>(n=43) | $\rho$ | 0.238 | 0.351 | 0.382 | 0.201 | -0.076 | 0.306 |
|  | *P*-value | 0.129 | 0.027* | 0.012** | 0.202 | 0.656 | 0.052 |
| **Total cholesterol**<br>(n=42) | $\rho$ | -0.122 | -0.014 | 0.293 | 0.085 | -0.148 | 0.052 |
|  | *P*-value | 0.445 | 0.932 | 0.060 | 0.599 | 0.390 | 0.748 |
| **HDL cholesterol**<br>(n=42) | $\rho$ | -0.118 | -0.077 | 0.138 | -0.036 | -0.223 | 0.178 |
|  | *P*-value | 0.461 | 0.640 | 0.384 | 0.825 | 0.190 | 0.265 |
| **Triglycerides**<br>(n=42) | $\rho$ | 0.206 | -0.051 | 0.157 | -0.039 | 0.033 | -0.221 |
|  | *P*-value | 0.197 | 0.759 | 0.321 | 0.810 | 0.847 | 0.164 |
| **Uric acid**<br>(n=43) | $\rho$ | 0.071 | -0.011 | -0.147 | 0.009 | 0.182 | -0.255 |
|  | *P*-value | 0.657 | 0.945 | 0.348 | 0.954 | 0.280 | 0.108 |
| **BNP**<br>(n=24) | $\rho$ | 0.004 | -0.231 | -0.235 | -0.148 | 0.481 | -0.053 |
|  | *P*-value | 0.986 | 0.288 | 0.269 | 0.500 | 0.024* | 0.809 |

Spearman's rho correlation coefficient ( $\rho$ ) and *P*-value are shown for each miRNA (\**P* < 0.05; \*\**P* < 0.05). BNP: brain natriuretic protein; CKD: chronic kidney disease; HDL: high-density lipoprotein; miRNA: microRNA.

### Circulating miR-324-3p and miR-199a-5p altered expression is associated with CKD progression and mortality, respectively

Patients included in the *discovery cohort* were followed up for an average of 70 months and data about the progression of CKD, cardiovascular events, number of hospitalizations, need for RRT, CVD, and other mortality causes were retrieved from the medical records (Table 9).

**Table 9.**
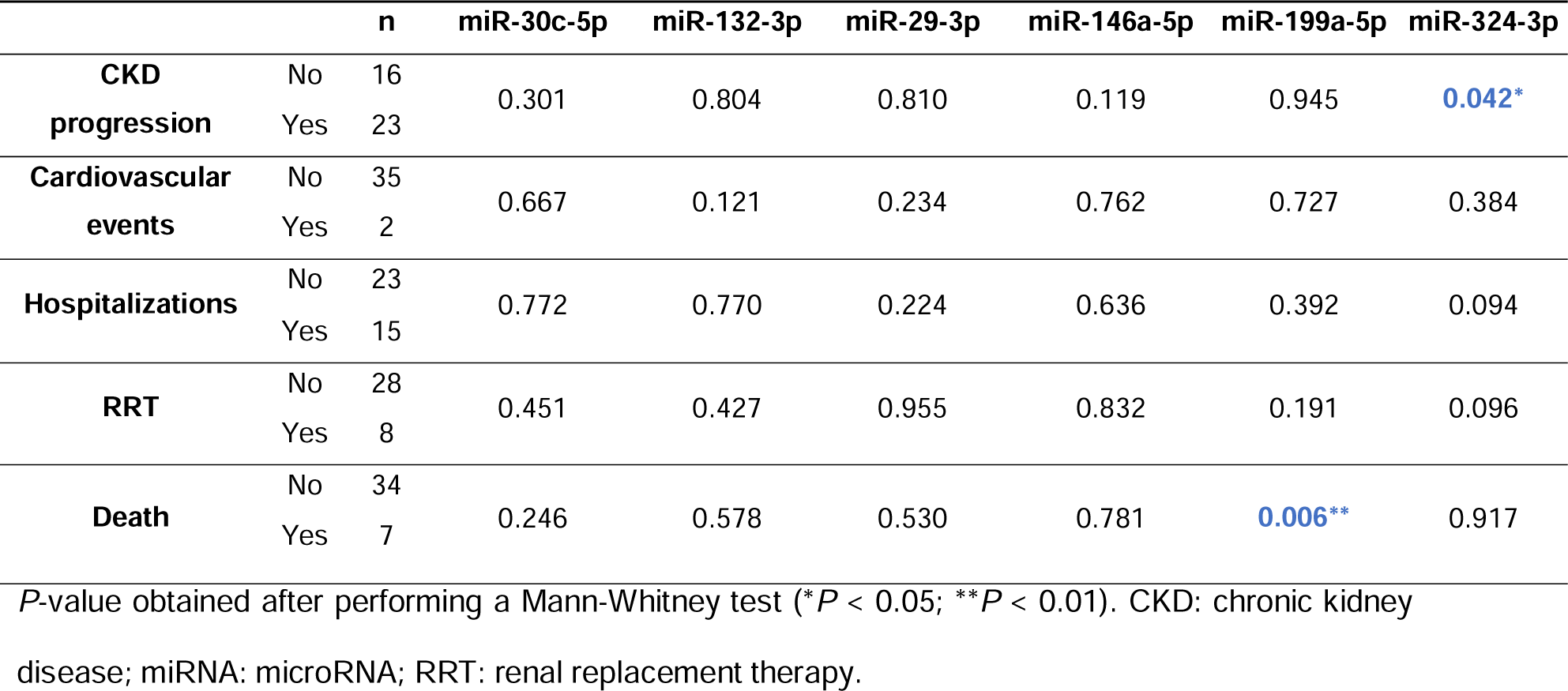
Comparative analysis of miRNA expression concerning CKD progression, cardiovascular events, hospitalizations, need for RRT, and mortality in a 5-year follow-up.

Throughout this period, 16 individuals progressed to a more advanced stage of CKD, and miR-324-3p was found to be downregulated in these patients in comparison with patients with non-progressive CKD (Figure 7a). Moreover, throughout the follow-up period, 7 of the CKD patients died, and statistical analysis showed that these patients exhibited elevated levels of miR-199a-5p (Figure 7b). Concerning the incidence of *de novo* cardiovascular events, only two patients with no previous history of CVD were identified, which is an insufficient number to allow significant statistical analysis. No significant differences were found in plasma miRNA expression regarding hospitalizations or the initiation of RRT in CKD patients of the *discovery cohort* during follow-up.

**Figure 7.**
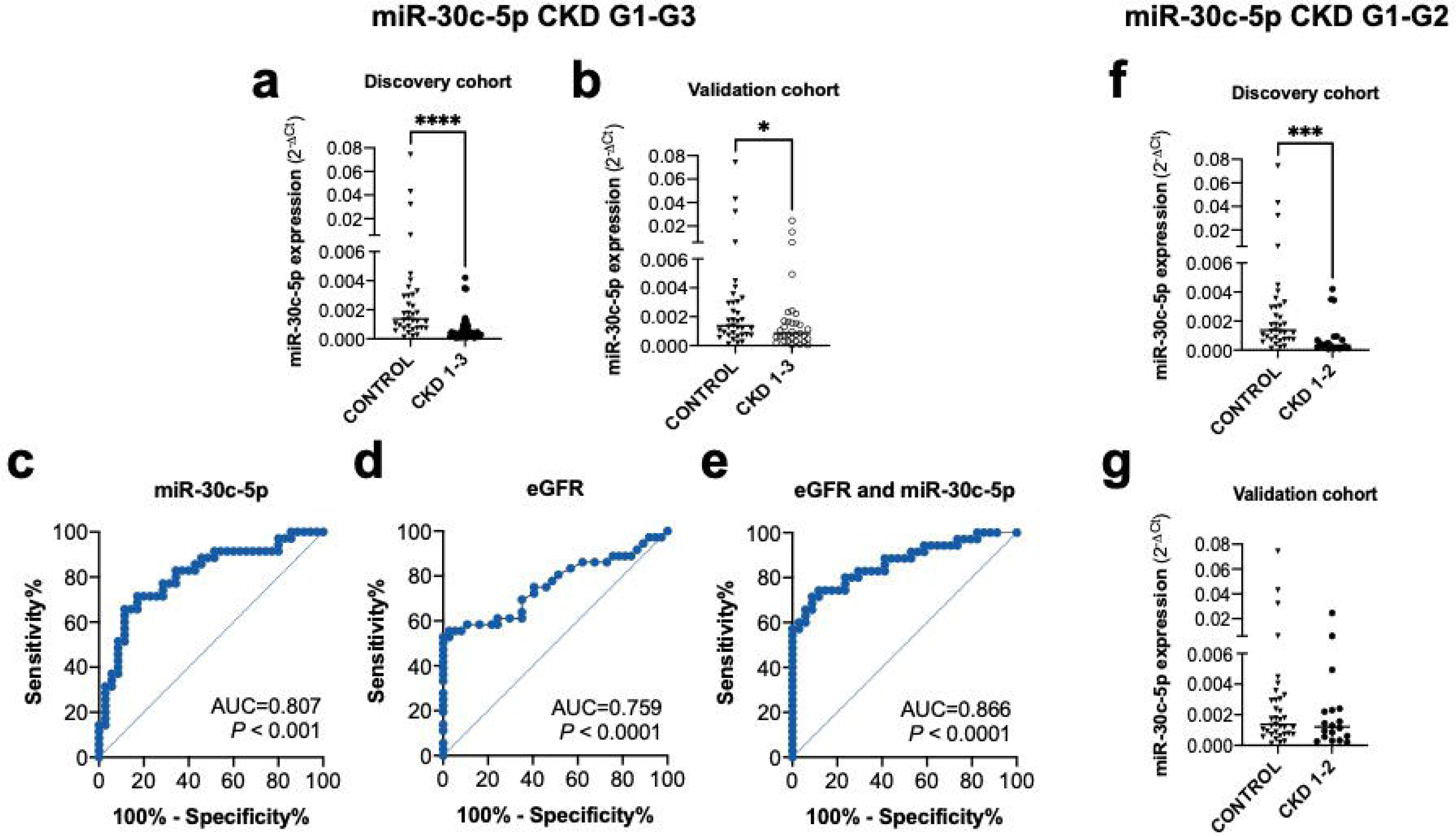
Circulating miR-324-3p and miR-199a-5p are associated with the progression of the disease and mortality in CKD patients, respectively. (a) MiR-324-3p expression is decreased in CKD patients who progressed to a worse CKD stage in a 5-year follow-up (n=16) in comparison with the ones that did not progress (n=23). (b) MiR-199a-5p expression is increased in CKD patients that died (n=7) within the 5-year follow-up period in comparison with the ones that did not (n=34). miRNA expression was quantified in plasma samples from the *discovery cohort* by real-time qRT-PCR (median; Mann Whitney test; \**P* < 0.05; \*\**P* < 0.01). CKD: chronic kidney disease; miRNA: microRNA.

**Figure 8.**
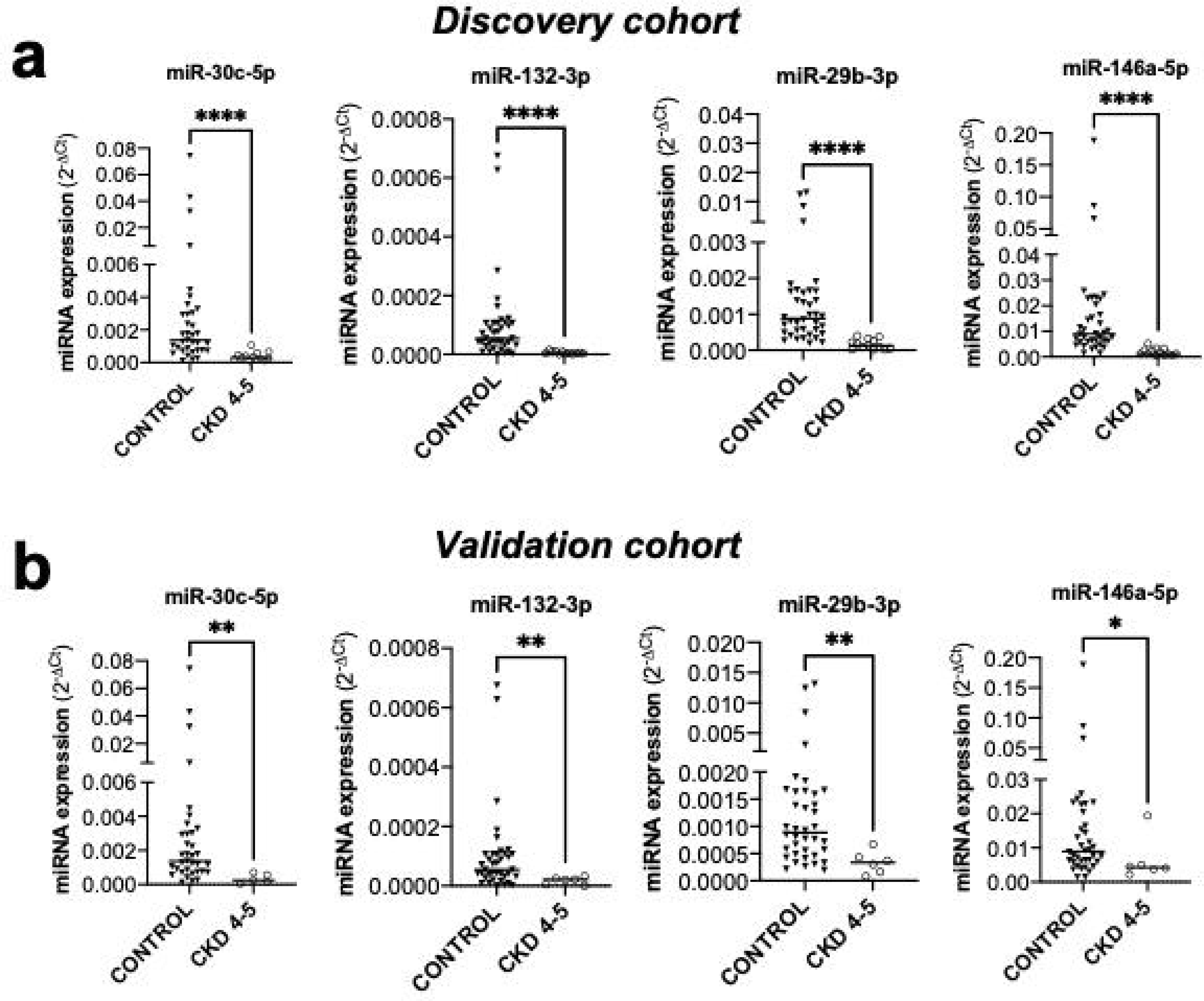
Summary figure describing the main findings of the current research article.

## Discussion

CKD represents a major health problem worldwide that poses many unmet clinical challenges, namely early diagnosis, prognosis, and therapeutic management options. The pathophysiology of CKD is very complex and poorly understood, and its diagnosis is frequently delayed due to the lack of symptoms in the early stages of the disease. In this context, circulating miRNAs emerge as promising candidates for biomarkers and therapeutic targets as they are accessible and exhibit high specificity and sensitivity to discriminate several pathological states ^20^.

Here, we evaluated the expression of a panel of miRNAs in plasma samples from CKD patients and its correlation with the clinical data at the time of sample collection and upon a 5-year follow-up time. The differential expression of miRNAs in CKD was assessed by comparison with its levels in plasma samples from healthy blood donors, with normal creatinine levels.

From the analyzed set of miRNAs, we observed a downregulation of miR-30c-5p and miR-132-3p in plasma samples from non-dialysis CKD patients compared to individuals with normal renal function. Such differential expression in CKD conditions was found to have good specificity and sensitivity for the diagnosis of the disease state, as supported by the AUC values of the ROC analysis (0.832 (*P* < 0.001) and 0.856, (*P* < 0.001) for miR-30c-5p and miR-132-3p, respectively). This fact, along with the significant increase observed in the AUC when combining the eGFR values with miR-30c-5p and miR-132-3p, compared to eGFR alone, suggests that these miRNAs could serve as alternative or complementary biomarkers for CKD diagnosis ^21^.

Remarkably, less than 5% of patients with early-stage CKD are aware of their condition, and the majority remain asymptomatic until they have reached a significant level of renal damage which impairs earlier interventions and is detrimental to disease prognosis ^3^. Aiming the investigation of these miRNAs as potential biomarkers for disease diagnosis, we evaluated its expression in CKD patients differentiated in early stages (G1-G2 and G1-G3, corresponding to mild and mild/moderate stages of disease, respectively) and in late disease stages (G4-G5). This analysis showed that miR-30c-5p was decreased in CKD patients since the early stages (G1-G2 only in the *discovery* cohort and G1-G3 in both the *discovery* and *validation* cohorts), showing a good discriminatory value from the healthy controls (AUC=0.807, *P* < 0.001 for G1-G3 analysis). Such relevance of miR-30c-5p in early CKD diagnosis was further sustained by logistic binary regression analysis where a statistically significant association between early CKD and miR-30c-5p was identified suggesting a protective effect of miR-30c-5p in CKD. Considering advanced CKD stages (G4 and G5), our investigation revealed that following up on the previous result, miR-30c-5p presented a progressively decreased expression from healthy to mild/moderate and severe CKD, and there was also a downregulation of miR132-3p, miR-29b-3p, and miR-146a-5p. The differential expression of some miRNAs in the early and late stages of the disease advocates that different pathways may be impacted throughout disease progression, with specific miRNAs being involved in each stage. This information may be of relevance not only for a better understanding of the biochemical pathways underlying disease progression but also for the design of specific strategies that can address CKD according to the disease stage. Of note is the fact that correlation analysis between miRNA expression and the patient’s clinical data showed that only miR-29b-3p and miR-132-3p were positively correlated with eGFR (as expected from its downregulation in CKD), while neither miR-30c-5p nor miR-146a-5p displayed a correlation. The absence of correlation between miR-30c-5p, miR-146a-5p and eGFR may be indicative of a putative role in CKD that is independent of eGFR, which may be groundbreaking for the establishment of new routes of investigation in this disease. In the case of miR-30c-5p, the fact that it was found to be significantly downregulated since early CKD stages, along with its ability to enhance CKD identification values in conjunction with eGFR in G1-G3, supports its potential as a biomarker of CKD in the early stages of the disease, when eGFR is still preserved.

The significant correlations identified between miRNA expression and other clinical data directly related to kidney function such as albumin (miR-29b-3p); P/C ratio (miR-199a-5p); CRP (miR-132-3p); vitamin D (miR-29b-3p) and calcium (miR-29b-3p and miR-30c-5p) allures that these miRNAs may exert different roles and target different metabolic pathways in CKD pathology that should be further investigated. Although there is currently very little data regarding the enrolment of miRNAs in kidney diseases in humans, our results are supported by previous studies conducted mostly *in vitro* or animal models of renal disorders. In fact, miR-30c-5p was shown to be decreased in several *in vitro* models of nephropathies, being involved in processes of oxidative stress, inflammation, extracellular matrix (ECM) deposition and epithelial-to-mesenchymal transition (EMT) ^22–26^. In contrast, a recent study showed upregulation of miR-30c-5p in tissue samples from IgAN patients within an Indian population ^27^. Such contradictory result highlights the intricate nature of miRNAs, namely the fact that they regulate several different pathways that may vary deeply among different populations and depend on disease specificities. In our cohorts, no significant differences were observed regarding the expression of miR-30c-5p among the different etiologies of CKD. Concerning miR-132-3p, it was found to be upregulated in renal tissue of a mouse model of ischemic acute kidney injury and to intensify the inflammatory response and oxidative stress in cultured proximal tubular cells ^28,29^. In contrast to these results, a recent study reported that this miRNA was downregulated in renal tissue in a sepsis-induced acute kidney injury mouse model ^30^ which is in line with our findings associating decreased expression of plasmatic miR-132-3p with worse kidney function and increased inflammation. The same correlations were found for miR-324-3p, suggesting the implication of both miRNAs in the modulation of pathways directly involved in renal function.

Regarding miR-29b-3p, some evidence indicates that its downregulation is associated with fibrosis in several organs, including the kidney ^31–35^. Likewise, *in vitro* and *in vivo* mice models of CKD reported a renal protective effect of miR-146a-5p, as its knockdown resulted in elevated tubular damage, inflammation, and fibrosis while its overexpression resulted in the opposite effect ^36–38^. Taking into consideration the fact that miR-29b-3p and miR-146 were decreased in CKD G4-G5, which are more prone to present renal fibrosis, the previously reported data supports a relevant anti-fibrotic activity that may be compromised in these sets of patients. This evidence may be of relevance for the establishment of stage-specific therapeutic approaches.

When addressing the kidney-heart interface, our study shows, for the first time to our knowledge, an association between plasmatic levels of miR-199a-5p and the presence of CVD in CKD patients, indicating a potential involvement in the pathophysiological communication between these two organs in CKD. This proposal is further supported by its positive correlation with P/C ratio; BNP, systolic blood pressure and left ventricular mass, all of them associated with an increased cardiovascular risk. Moreover, miR-199a also showed a negative correlation with ejection fraction. Although this parameter is not a defining factor for the existence of heart disease, patients with heart failure and reduced ejection fraction exhibit a more typical and more severe phenotype.

Previous studies documented independent roles of this miRNA in both the renal and cardiac system but its putative role in the kidney-heart communication was not addressed. It was shown to play a role in the development of cardiac hypertrophy ^39–41^ and to be increased *in vitro* and *in vivo* kidney disease models targeting klotho, a protein with anti-aging and antioxidant characteristics whose deficiency has been associated with the development and progression of CKD ^42–44^. Regarding miR-324-3p, although our results show decreased expression in CKD patients that progressed into more advanced stages, most documented studies report a pro-fibrotic role of this miRNA in renal disease ^45–47^. Such apparently conflicting results may indicate that miR-324-3p can play a role in the regulation of CKD progression that is not directly associated with fibrosis.

During the follow-up period, 23 patients experienced CKD progression, 2 patients suffered cardiovascular events, 23 were hospitalized, 8 needed RRT and 7 died. Taking into consideration the low number of events in some of these variables, our analysis was hampered by limited statistical power. An increase in the number of patients enrolled and longer follow-up periods would help to clarify the predictive power of the identified circulating miRNAs for forecasting the mentioned events.

Summing up, our results provide compelling evidence that circulating miR-30c-5p and miR-132-3p may serve as plasmatic biomarkers for CKD diagnosis. In particular, miR-30c-5p has shown significant potential as a biomarker for early CKD diagnosis and prediction, being enrolled throughout all the stages of the disease, while miR-132-3p, miR-29b-3p, and miR-146a-5p were shown to be decreased only in advanced CKD states. When considering major CKD outcomes, miR-199a-5p was associated with the presence of CVD and mortality, while miR-324-3p downregulation was related to CKD progression.

Overall, these findings identify new miRNAs involved in the onset and progression of CKD and pinpoint miR-199a-5p as a relevant player in the kidney-heart pathophysiological crosstalk. These results may pave the way for the discovery of the long-searched biomarkers and therapeutic targets for CKD progression and CVD outcomes, with a high potential for translation into clinical practice. Further studies involving larger and more diverse cohorts of patients across multiple centers, and mechanistic studies are essential for fully understanding the enrolment of these miRNAs in CKD.

## Acknowledgements

The authors acknowledge the Nephrology Service of CHUSJ do Porto, for assisting with the sample collection of the patients included in this study, and the Immunotherapy Service of CHUSJ, for kindly providing the samples of healthy donors. The authors thank Professor Pedro Oliveira, from the Institute of Biomedical Sciences Abel Salazar (ICBAS), University of Porto for the kind assistance with statistical analysis.

All authors have read the journal’s policy on disclosure of potential conflicts of interest and declare no conflict of interest.

## Data sharing statement

All data is included in the manuscript and supporting information.

## Author contributions: CRediT

Olalla Ramil-Gómez: data curation, formal analysis, investigation, methodology, software, supervision and, writing – original draft and writing – review and editing. Ana Cerqueira: data curation, investigation, resources, validation and writing – review and editing. Sara Reis Moura: methodology, software, Writing – review and editing. Eduarda Carvalho: investigation and writing – review and editing. Núria Paulo: data curation, resources, validation and writing – review and editing. Isabel Brandão: investigation and writing – review and editing. Janete Santos: data curation and resources. Patricia Quaranta:: software and formal analysis. Maria João Sousa: resources. Ana Pinho: software and formal analysis. Hugo Diniz: resources, supervision, validation and writing – review and editing. Maria Inês Almeida: data curation, investigation, methodology, resources, supervision, validation and writing – review and editing. Manuel Pestana: conceptualization, funding acquisition, supervision and writing – review and editing. Inês Soares Alencastre: conceptualization, data curation, funding acquisition, investigation, project administration, resources, supervision, validation, visualization, writing – original draft and writing – review and editing.

---

**Figure S1.**
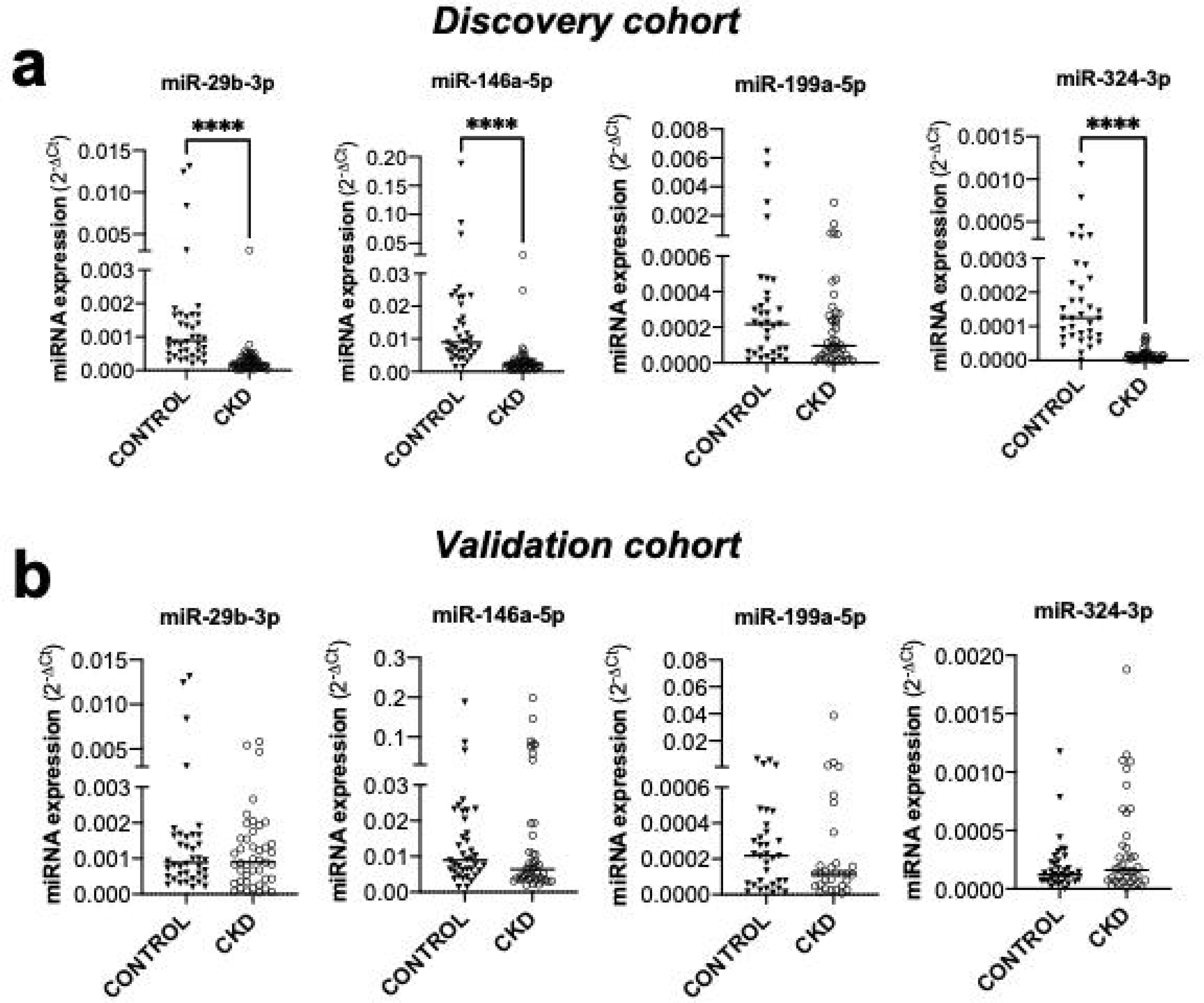
Expression of circulating miR-29b-3p, miR-146a-5p, miR-199a-5p, and miR-324-3p in CKD patients *vs* healthy controls. miRNA expression was assessed in plasma samples from the control group (n=38) and CKD patients from (a) the *discovery* (n=52) and (b) *validation* (n=43) *cohorts* by real-time qRT-PCR. (Median; Mann-Whitney test, \*\*\*\**P* < 0.0001). CKD: chronic kidney disease; miRNA: microRNA.

